# Single-cell isoform RNA sequencing (ScISOr-Seq) across thousands of cells reveals isoforms of cerebellar cell types

**DOI:** 10.1101/364950

**Authors:** Ishaan Gupta, Paul G Collier, Bettina Haase, Ahmed Mahfouz, Anoushka Joglekar, Taylor Floyd, Frank Koopmans, Ben Barres, August B Smit, Steven Sloan, Wenjie Luo, Olivier Fedrigo, M Elizabeth Ross, Hagen U Tilgner

**Affiliations:** Brain and Mind Research Institute and Center for Neurogenetics, Weill Cornell Medicine, 413 east 69^th^ Street, New York, NY 10021; The Rockefeller University, 1230 York Avenue, New York, NY, 10065; Leiden Computational Biology Center, Leiden University Medical Center, Albinusdreef 2, 2333 ZA Leiden; Delft bioinformatics Lab, Delft University of Technology, van Mourik Broekmanweg 6, 2628 XE Delft; Dept. Molecular and Celullar Neurobiology, Center for Neurogenomics and Cognitive Research, Amsterdam Neuroscience, VU University, Amsterdam, The Netherlands; Department of Neurobiology, Stanford University, 299 Campus Dr, Stanford, CA 94305-5125; Brain and Mind Research Institute and Appel Alzheimer’s research institute, Weill Cornell Medicine

## Abstract

Full-length isoform sequencing has advanced our knowledge of isoform biology^1–11^. However, apart from applying full-length isoform sequencing to very few single cells^12,13^, isoform sequencing has been limited to bulk tissue, cell lines, or sorted cells. Single splicing events have been described for <=200 single cells with great statistical success^14,15^, but these methods do not describe full-length mRNAs. Single cell short-read 3’ sequencing has allowed identification of many cell sub-types^16–23^, but full-length isoforms for these cell types have not been profiled. Using our new method of single-cell-isoform-RNA-sequencing (ScISOr-Seq) we determine isoform-expression in thousands of individual cells from a heterogeneous bulk tissue (cerebellum), without specific antibody-fluorescence activated cell sorting. We elucidate isoform usage in high-level cell types such as neurons, astrocytes and microglia and finer sub-types, such as Purkinje cells and Granule cells, including the combination patterns of distant splice sites^6–9,24,25^, which for individual molecules requires long reads. We produce an enhanced genome annotation revealing cell-type specific expression of known and 16,872 novel (with respect to mouse Gencode version 10) isoforms (see isoformatlas.com).

ScISOr-Seq describes isoforms from >1,000 single cells from bulk tissue without cell sorting by leveraging two technologies in three steps: In step one, we employ microfluidics to produce amplified full-length cDNAs barcoded for their cell of origin. This cDNA is split into two pools: one pool for 3’ sequencing to measure gene expression (step 2) and another pool for long-read sequencing and isoform expression (step 3). In step two, short-read 3’-sequencing provides molecular counts for each gene and cell, which allows clustering cells and assigning a cell type using cell-type specific markers. In step three, an aliquot of the same cDNAs (each barcoded for the individual cell of origin) is sequenced using Pacific Biosciences (“PacBio”)^1,2,4,5,26^ or Oxford Nanopore^3^. Since these long reads carry the single-cell barcodes identified in step two, one can determine the individual cell from which each long read originates. Since most single cells are assigned to a named cluster, we can also assign the cell’s cluster name (e.g. “Purkinje cell” or “astrocyte”) to the long read in question (Fig 1A) – without losing the cell of origin of each long read.

## Results

### Detection of cell types

We apply ScISOr-Seq to describe cell-type specific isoforms in mouse cerebellum at postnatal day 1 (P1). We sequence a mean of 17,885 reads per cell (as given by 10xGenomics’ summary statistics). After filtering cells and considering only reads confidently mapped to genes, we have 3,875 unique molecular identifiers (UMIs) and 1,448 genes per cell during 3’end sequencing. In step 2, 6,627 cells were clustered into 17 groups (Figs. 1B,D). High expression of well-established cell-type specific markers identifies many clusters as cell types: High expression of *Pdgfra*, *Olig1* and *Olig2* identified a cluster of oligodendrocyte precursors (OPCs, Fig 1B,C). *Clu* and *Apoe* identified two clusters of astrocytes and *Gdf10*^27,28^ identified a cluster of Bergmann Glia (BG). We also identified three large clusters of neuronal subtypes: namely (i) cells with high expression of *Neurod1* and *Ccdn2*, which we refer to as external granular layer (EGL) cells in several stages of differentiation. These give rise primarily to granule neurons that migrate into the internal granular layer (IGL) over the first weeks of mouse postnatal development; (ii) Purkinje-like (marked by *Pcp4, Gad1* and *Gad2*) in the Purkinje cell layer (PCL) and (iii) other neurons known to be present in the deep cerebellar nuclei and internal granular layer (collectively referred to as “IGL” heareafter), in which cells display high expression of *Pnoc*, *Snhg11*, *Tcf7l2*, *Gad1*, *Gad2* and *Lhx9*. This cluster was clearly composed of at least two clusters: One expressing *Gad1* and *Gad2* and the other expressing *Lhx9* and *Tcf7l2* (Figure 1B). These cell-type specific expression patterns exhibit specific anatomical localization within the developing cerebellum (Figure 1C, Allen Brain Atlas). Three further clusters expressed genes associated with neural progenitor cells: one expressing *Ccnd2* (which is highly expressed in the postnatal EGL), another *Atoh1* (glutamatergic neuron precursors from the rhombic lip and EGL) and the third *Ptf1a* (GABAergic neuron precursors from the ventricular zone) (Fig 1B). We could also identify other cell populations, such as microglia, which highly expressed well-known myeloid-associated genes (e.g. *C1qa*, *C1qb*, *C1qc* and *Tmem119)*. Alongside neuroglial subtypes and progenitors, we furthermore identified clusters expressing genes specific to endothelial and circulatory-system cells. In summary, our clustering recapitulates a large proportion of cell types classically observed in P1 cerebellum. Overall EGL, IGL cells and astrocytic cells were the largest cluster and blood cells the smallest. Detected reads, short-read UMIs and genes per cell showed slight differences between cell types but were of similar orders of magnitude. Consistent with their large size, Purkinje cells had the highest number of read, UMI and gene counts, while blood cells had the lowest gene count.

**Figure 1:**
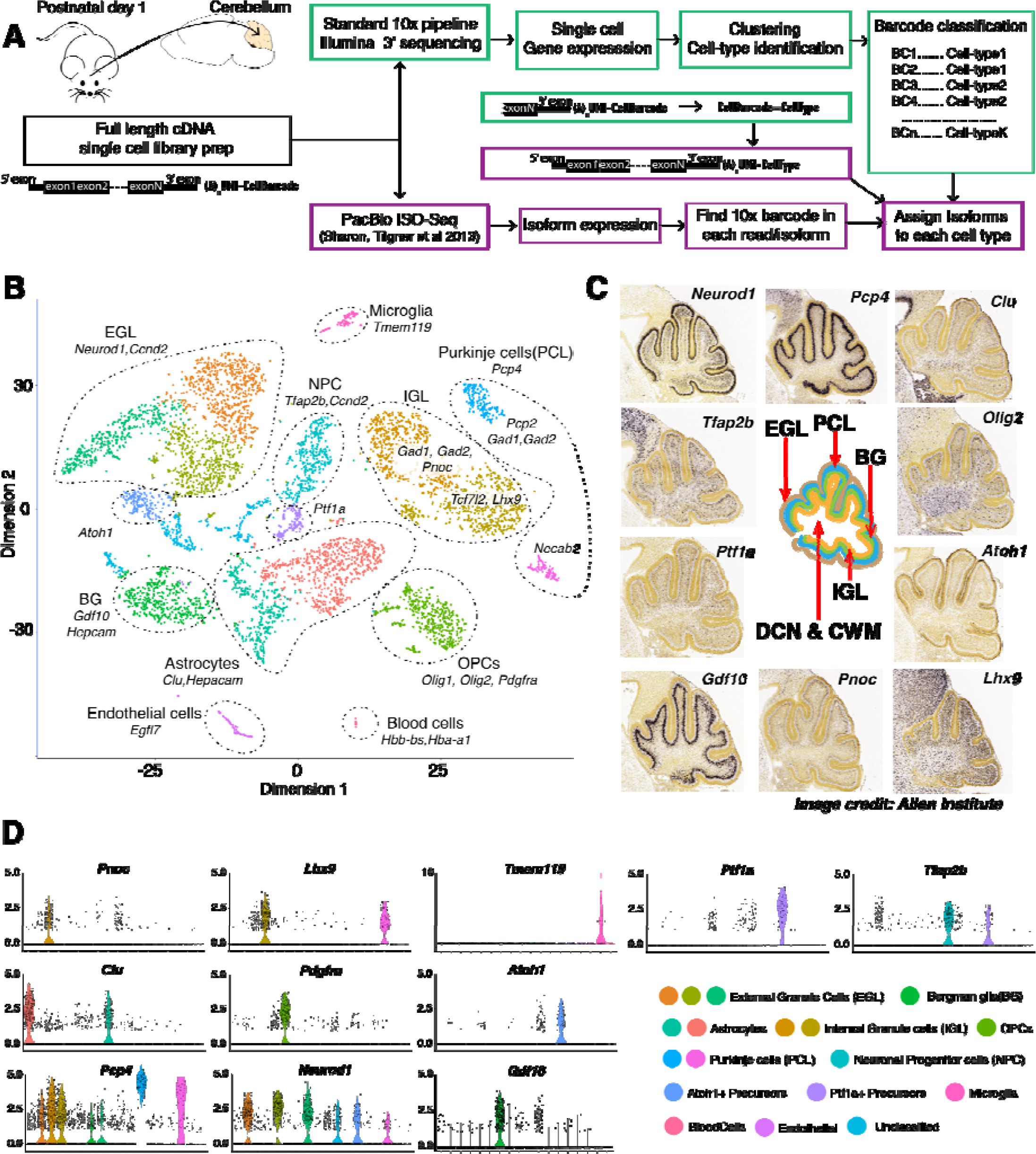
(A) Outline of our ScISOr-Seq approach. (B) TSNE-plot depicting cell clusters, marker genes and names given to clusters, including: Bergman glia (BG), External granule cell layer neurons (EGL), Internal granule cell layer and other neurons in the interior of the cerebellum (IGL), two clusters of Purkinje cell layer neurons (PCL), oligodendrocyte progenitor cells (OPCs), Atoh1+ neuronal progenitors, Ptf1a+ neuronal progenitors and other neuronal progenitors (NPCs) (C) In-situ hybridization images from the Allen Brain Atlas depicting expression of marker genes in specific layers. (D) Expression patterns of selected marker genes across cell types.

### Reliability and replication of cell-type detection

Sequencing of a second replicate (rep2) and within-replicate analysis showed that most distinct clusters were highly dissimilar to any other clusters in the same replicate. To assess stability of clusters, we tripled Illumina sequencing depth for rep2. In all clusters (with one exception) based on shallower sequencing depth, 95-100% of cells were still attributed to the same cluster, even with three-fold deeper sequencing. Analysis of comparability of marker gene between clusters of the two replicates using the Jaccard index identified highly similar clusters with one exception: The smallest cluster (blood cells) in replicate 1 (rep1), was missing in rep2. Cell-type abundance was reproducible between replicates and highly correlated (Pearson correlation = 0.91, correlation-test p-value = 4.5 ×10^−5^).

### Detection of single-cell barcodes in full-length cDNAs

We then employed 850ng of full-length cDNAs, tagged for their cell of origin, for isoform sequencing to generate ~5.2 million PacBio circular consensus reads (“CCS”). These CCS showed mean full passes per SMRT cell of 16-34 and thus favor a lower error rate compared to earlier ISO-Seq publications^1,4^. Since cellular barcodes are located close to the polyA-tail, we first searched for polyA-tails. Aiming at detecting polyT-sequences, even with a hypothetical 10% error rate, we located the first nine consecutive Ts (“T9”) in the first 200bp of each read and of its reverse complement. 61.6% of CCS contained such a T9, broadly consistent with our previous estimation (67%)^1,4^. Reads with and without T9s showed similar lengths, apart from CCS <=200bp accumulating in non-T9 CCS. 1.4% of T9-CCS had a T9 in the read start and the complement’s start. These may include chimeras, which were removed from further analysis, introduced during reverse transcription, PCR or blunt-end PacBio library preparation. In total, for 58.0% (compared to 74.0% for 10x-3’seq) of the polyA-tail-containing CCS, we identified a perfect-match 16mer cellular barcode (each corresponding to one of the 6,627 single cells) and therefore the exact single cell, in which the RNA isoform was transcribed. As a theoretical foundation, we determined for all 6,627 barcodes, the minimal editing distance to any other barcode: For 92.7% of barcodes, this minimal (“Levenshtein”) distance was 3 or greater, and for the remaining barcodes it was two. This shows that for most barcodes, there is one specific error pattern of three errors that would lead to a wrongly identified cell. However, in most cases three random errors would only discard the read because none of the 6,627 known barcodes is detected. Both experiments and simulations show that our single-cell barcode-detection procedure is extremely specific. Overall, we detected a median of 270 long reads, 260 UMIs and 129 genes per single cell. 3.8% of UMIs are observed twice, with a theoretical prediction of 3.4% (Methods). 99.3% (6,581/6,627) of clustered cells were detected with CCS (Figure 2A-D). 97.4% (6,459/6,627) of clustered cells had >100 CCS (Figure 2D). Detected short-read and long-read UMIs per single cell correlated highly (Pearson correlation = 0.95, correlation-test p < 2.2 × 10^−16^, Figure 2E). Long-read statistics (reads, UMIs, genes, Figure 2F-H) per cell cluster mirrored those in short-reads, with lower long-read numbers.

**Figure 2:**
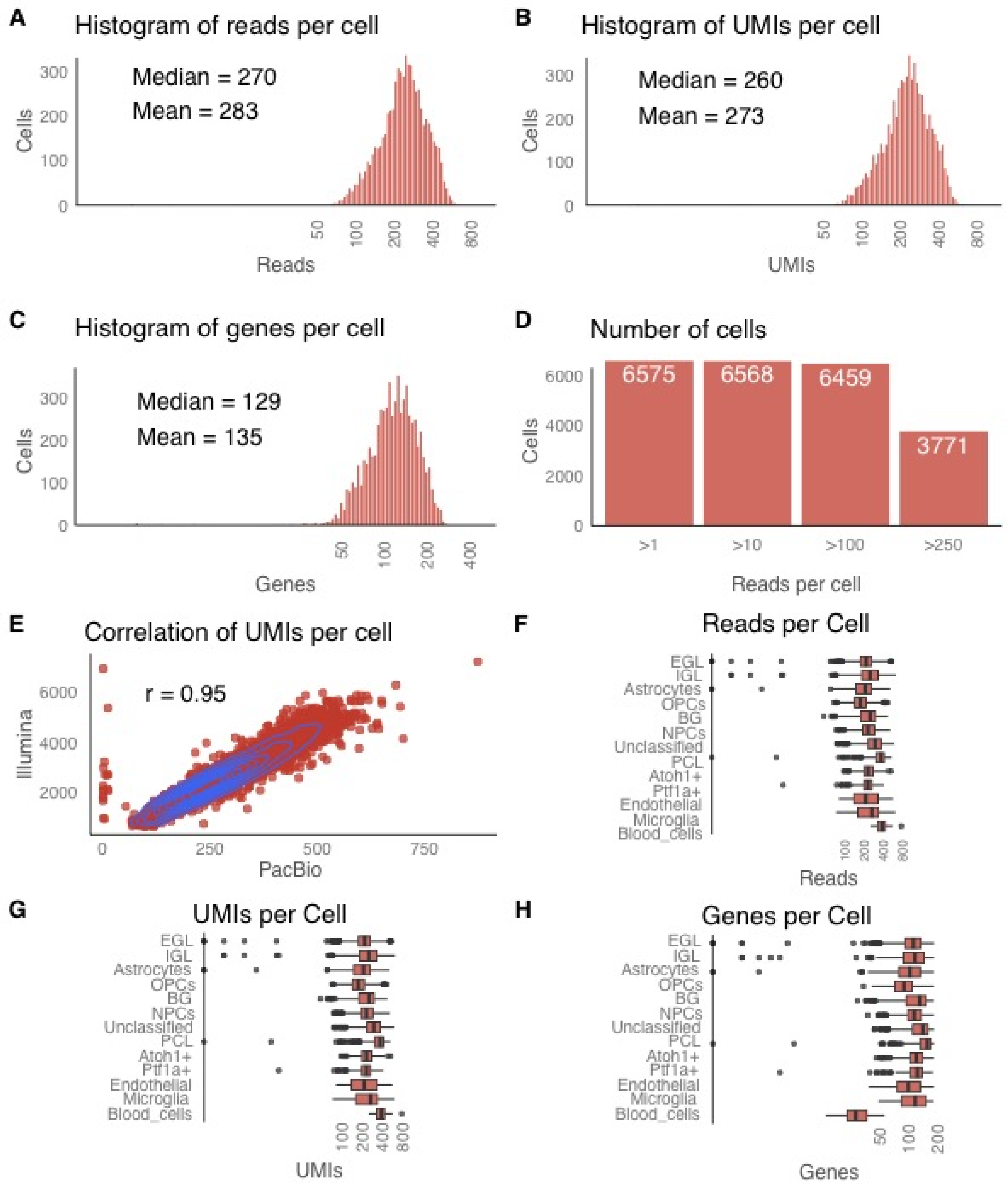
Long-read statistics. Distribution of (A) long reads (B) long-read UMIs and (C) genes per cell. (D) Number of cells >1,>10,>100,>250 long-reads (E) Dotplot and correlation between long-read UMIs and short-read UMIs per cell. Distribution of (F) long reads (G) long-read UMIs and (H) long-read detected genes per cell.

### ScISOr-Seq using Nanopore sequencing

Using 1µg of barcoded cDNA on a Nanopore MinIon, we searched for cellular barcodes in 2.3 million Nanopore reads^29^. We found lower relative numbers of Nanopore 1D reads with a T9, supposedly due to problems with homopolymers in Nanopore data^29^. However, ~31.4% (1D) and ~35.2% (passed 1D^2^) of Nanopore reads have a 30bp window with >=25 Ts. Although the variation from the expected position in Nanopore reads is larger than for CCS (90bp vs. 3bp), accumulation around the expected position is observed and exact barcode matches reveal unique barcodes in 6.0% (43,948/732,590, 1D) and 32.7% (9,454/28,931, 1D^2^). Therefore, we can expect ~50,000 cluster-specific long reads per MinIon flow cell. With each current MinIon flow cell requiring 1µg of cDNA, further PCR (with associated biases) is needed to carry out large-scale ScISOr-Seq on Nanopore, whereas the employed 16-cycle PCR is sufficient to run 20-50 SMRTcells on PacBio yielding up to 5 million long reads assigned to single cells.

### A cell-type resolved isoform annotation

We aligned PacBio CCS to the mouse genome^30^(version mm10) using STAR^31^and carried out mapping quality control as previously performed^1,4,6^. We analyzed novel isoforms with respect to mouse Gencode version 10, as outlined previously^1,6,32^ to produce a long-read enhanced and cell-type resolved annotation. We considered 10,691 unique novel (with respect to mouse Gencode version 10) isoforms that affected 4,859 genes. For these isoforms, we required all splice sites to be known in Gencode^33^ (version 10) and each junction and internal exon to be either annotated or observed at least twice in ScISOr-Seq. The unique novel isoforms contain new exon-exon junctions linking previously known splice sites, such as the skipping of exons annotated as constitutive. Artifacts in next-generation sequencing have been demonstrated^34^. To assess whether the long-range 16-cycle PCR in ScISOr-Seq generates chimeric transcripts, we obtained 164 million 150bp-paired-end reads on bulk RNA from P1 cerebella only employing a 6-cycle short-range PCR after RNA fragmentation. Based on this experiment, we confirmed 91.6-97.6% of the novel ScISOr-Seq junctions across different cell types (Figure 3A). To reduce the influence of PCR artifacts on the enhanced annotation to a minimum and to allow for adding lowly expressed transcripts, we generated a final enhanced cell-type resolved annotation with strong 6-cycle-PCR short-read support. In this enhanced annotation, for each added isoform, each intron and internal exon was required to be annotated in Gencode or to be supported by two or more 6-cycle-PCR short reads, resulting in 16,872 isoforms for 6,927 genes (Figure 3B). For each of these isoforms we know the single cell of origin and therefore the cell type that produced this isoform. 42.8% (7,219/16,872) employed at least one splice site not annotated in Gencode. With respect to the UCSC^35^and RefSeq^36^annotations, 94.0% and 70.9% respectively of added isoforms were novel We performed ScISOr-Seq for rep2, albeit at a lower sequencing depth (6 SMRT cells, compared to 23 for rep1). New rep2-isoforms replicated in rep1 in 65.7% (microglia) to 76.2% (NPCs)(Figure 3C) of the cases (irrespectively of the cell type they were observed in rep1). Given replication of an isoform in any cell type, cell-type specific replication of a rep2-isoform in the same cell type in rep1 reached 70-80% in larger clusters, but lower percentages in smaller clusters with dramatically fewer long reads (Figure 3D). To validate the correct calling of the individual cell of origin for each isoform, we performed immunopanning to specifically isolate microglia in P1 cerebella followed by short-read RNAseq. This data was compared to all isoforms originating from a single microglial cell (and then to isoforms of single cells belonging to other cell types). This confirmed the microglial origin of long-read junctions exclusively observed in microglial single-cell long reads as compared to junctions observed exclusively in non-microglial single-cell long reads (Figure 3E). Similarly, immunopanning for astrocytes, Bergmann glia (both marked by *Hepacam)* and OPCs (which are known to be enriched in *Hepacam-*sorting) and short-read sequencing showed the highest coverage for junctions observed exclusively in astrocyte, Bergmann Glia and OPC ScISOr-Seq isoforms This was more pronounced for junctions observed three or more times in ScISOr-Seq data in one cell type. These data suggest that junctions observed only in astrocytes, Bergmann Glia and OPCs are also expressed at a lower level in other cell types originating from the same stem cell.

**Figure 3:**
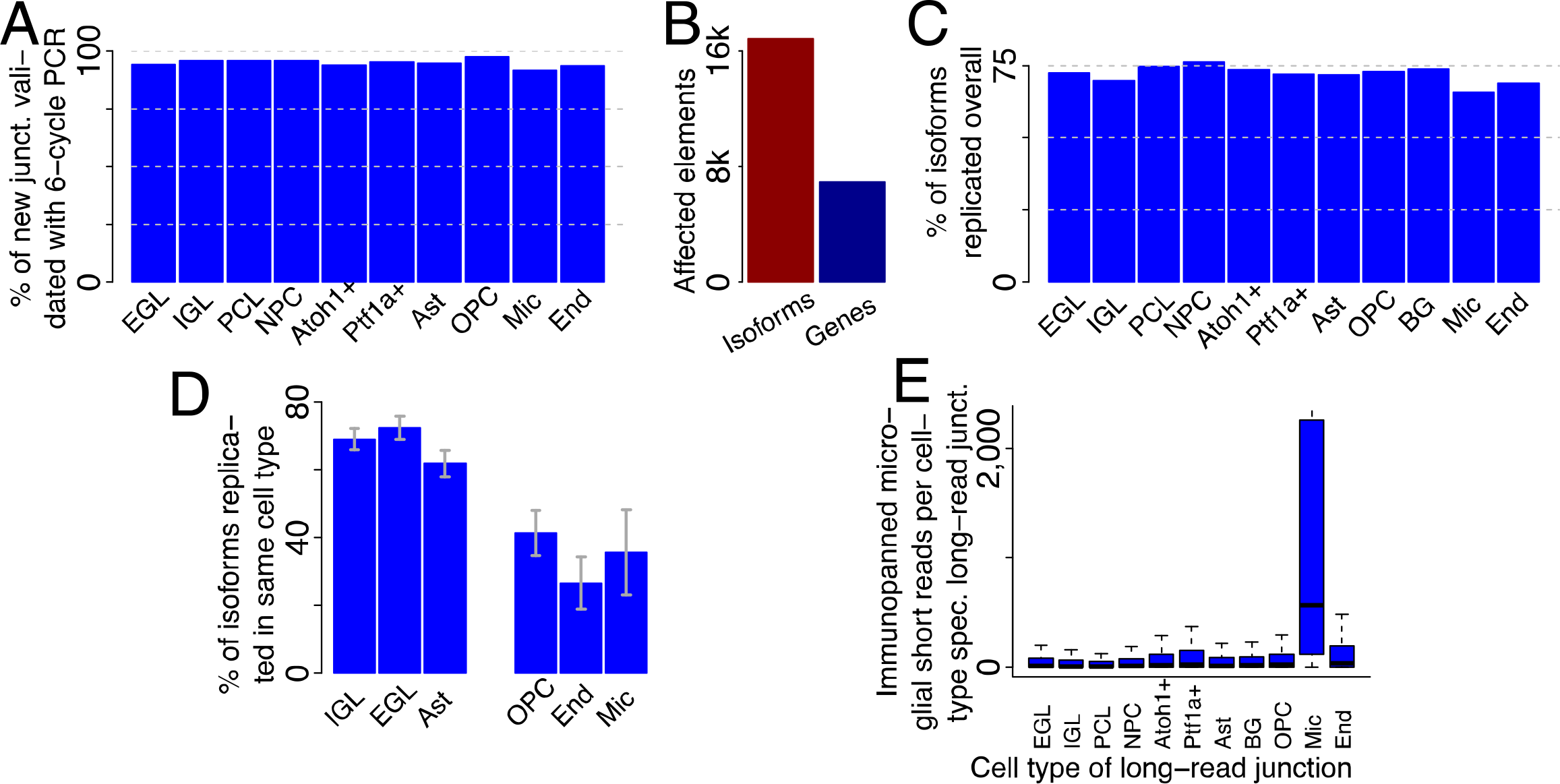
An enhanced and cell-type specific annotation. (A) Percentage of long-read derived junctions that could be validated using low-cycle PCR from bulk P1 cerebellum. (B) Number of isoforms added to the annotation and number of affected genes. (C) Percentage of complete unique isoforms from replicate 2 that could also be observed in replicate 1 (<u>in any cell type</u>) broken up by cell type of origin from replicate 1. (D) Percentage of complete unique replicated isoforms from replicate 2 that could also be observed in replicate 1 (<u>in the same cell type</u>) broken up by cell type of origin from replicate 1 (E) Distributions of coverage with microglial short reads for introns in the enhanced annotation that were exclusively observed in one cell type (indicated by name under the x-axis).

### Database of cell-type specific isoform expression in the cerebellum

We first looked at alternative splicing in *Tpm1* gene that is expressed across multiple cell types and is known to have extensive alternative splicing according to Gencode^33^, UCSC^35^ and RefSeq^36^ annotations. This gene contains five alternatively spliced blocks of exons namely AS1-AS5 (observable in >=3 reads). AS1 and AS5 represent alternatively spliced blocks of single or multiple 5’ and 3’ exons along with the associated untranslated regions while AS2-AS4 represent single alternatively spliced exons within the coding region of the gene. We observed 4 novel isoforms (Figure 4, black and bold) of *Tpm1* as compared to the observed 21 isoforms according to the Gencode annotation. Out of these “Novel Isoform 4”, where AS4 is spliced out while other alternatively spliced exons are included, was the major isoform expressed in Astrocytes with 10 UMIs. Out of the annotated transcripts, only the OPCs express the ENSMUST00000113685.9 (Figure 4, red) transcript with 15 UMIs which was also the most abundantly expressed isoform in OPCs. Other annotated transcripts ENSMUST00000113686.7 (Figure 4, orange) and ENSMUST00000113690.7 (Figure 4, green) were the most abundant isoforms in EGL and IGL respectively. In order to make this data accessible to the research community, we have created a fully searchable database (see isoformatlas.com) of isoforms for every gene showing their cell type of origin (as shown in Figure 4) and their single-cell of origin projected onto the TSNE-plot shown in Figure 1B.

**Figure 4:**
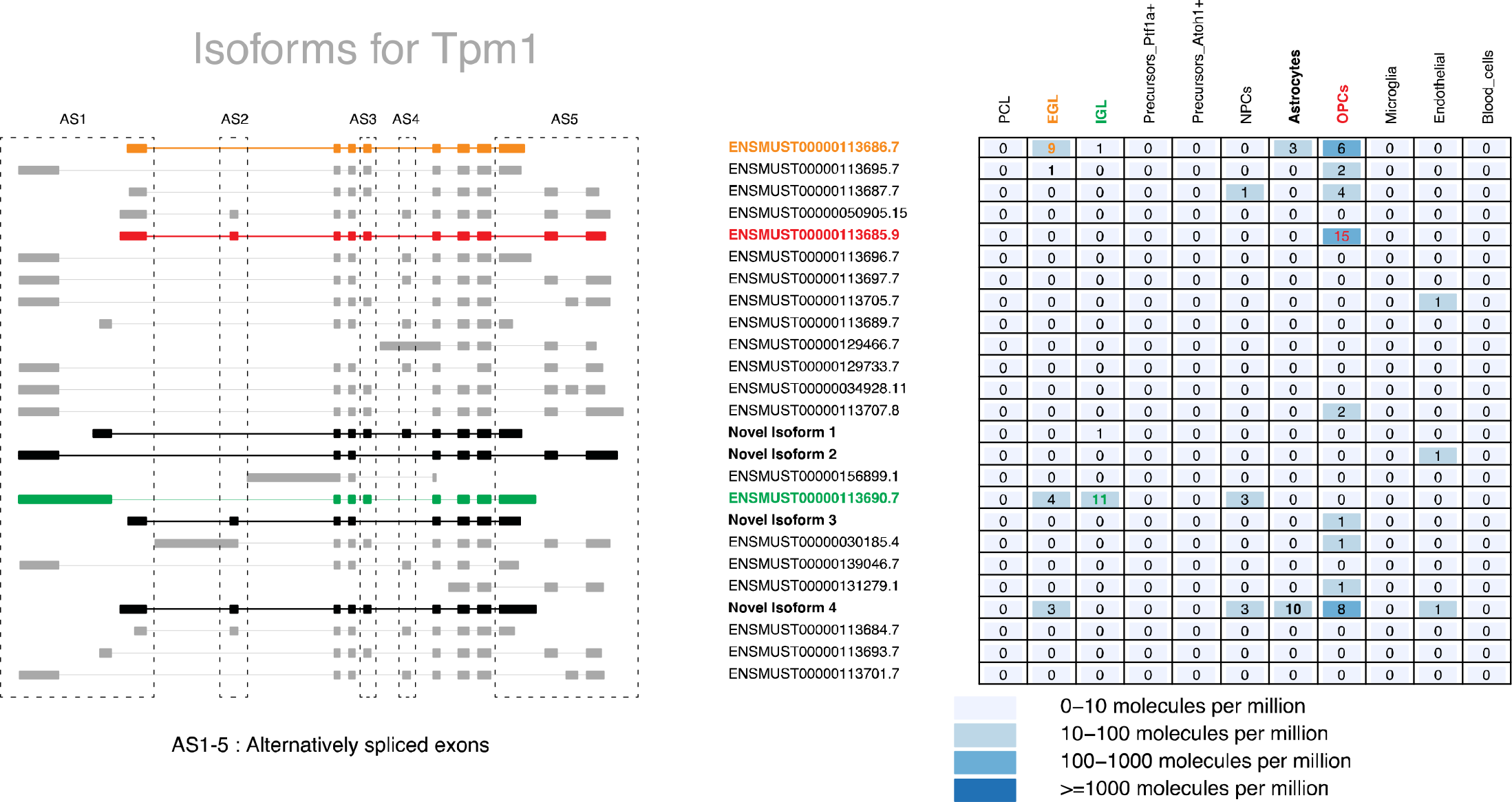
Single gene view for the *Tpm1* gene (from isoformatlas.com), Left: Isoforms of the gene, where each exon is a rectangular block joined by an intervening line representing introns. Alternatively spliced exon blocks identified by AS1-5 are enclosed by dashed-lined boxes Right: Table representing the distribution of UMI counts per isoform (in rows) and cell type of origin as identified in Figure 1 (in columns). Major isoforms in each cell type are colored orange (EGL), red (OPCs) and green (IGL). Novel isoforms are colored in black.

## Discussion

Brain disorders (e.g. Alzheimer’s disease) are highly associated with risk genes including *MAPT*and *APOE*. Interestingly, these genes are expressed in multiple cell (sub-)types. Therefore, cell-type specific isoform expression is critical and may decipher the action of disease-associated SNPs. Here, we (i) describe isoform expression across heterogeneous cell types and (ii) enhance genome annotation with cell-type specific isoform expression. A drawback is that employing multiple deeply sequenced replicates is for now very expensive with long-reads, making precise quantification of abundance changes between cell types as outlined in rMATS^37^more difficult. However, our full-length RNAs from single cells cover all single nucleotide polymorphisms in the coding region of mature RNA and may help attributing single cells to a specific individual^38^in pooled samples.

## Acknowledgements

This work was supported by start-up funds (Weill Cornell Medicine) and a Leon Levy Fellowship in Neuroscience to HUT. This work used the Genomics Resources Core Facility and owes special thanks to Dr. Jenny Xiang and Angela Wan.

## Author contributions

Devised the experiments: PGC, IG, SS, HUT. Performed experiments: PGC, BH, IG, SS, OF, WL; Devised analysis: IG, AS, HUT. Performed analysis: IG, AM, AJ, TF, FK, HUT; Discussed and interpreted results throughout the project: all authors; Wrote the paper: IG, HUT with inputs from all authors. Supervised research: BB, MER, HUT.

